# QSM Reconstruction Challenge 2.0: a realistic in silico head phantom for MRI data simulation and evaluation of susceptibility mapping procedures

**DOI:** 10.1101/2020.09.29.316836

**Authors:** José P. Marques, Jakob Meineke, Carlos Milovic, Berkin Bilgic, Kwok-Shing Chan, Renaud Hedouin, Wietske van der Zwaag, Christian Langkammer, Ferdinand Schweser

## Abstract

**Purpose:** To create a realistic *in-silico* head phantom for the second Quantitative Susceptibility Mapping (QSM) Reconstruction Challenge and for future evaluations of processing algorithms for QSM.

**Methods:** We created a digital whole-head tissue property phantom by segmenting and post-processing high-resolution (0.64mm isotropic), multi-parametric MRI data acquired at 7T from a healthy volunteer. We simulated the steady-state magnetization at 7T using a Bloch simulator and mimicked a Cartesian sampling scheme through Fourier-based processing. Computer code for generating the phantom and performing the MR simulation was designed to facilitate flexible modifications of the phantom in the future, such as the inclusion of pathologies, as well as the simulation of a wide range of acquisition protocols. Specifically, the following parameters and effects were implemented: repetition time and echo time, voxel size, background fields, and RF phase biases. Diffusion weighted imaging phantom data is provided allowing future investigations of tissue microstructure effects in phase and QSM algorithms.

**Results:** The brain-part of the phantom featured realistic morphology with spatial variations in relaxation and susceptibility values similar to the in vivo setting. We demonstrated some of the phantom’s properties, including the possibility of generating phase data with non-linear evolution over echo time due to partial volume effects or complex distributions of frequency shifts within the voxel.

**Conclusion:** The presented phantom and computer programs are publicly available and may serve as a ground truth in future assessments of the faithfulness of quantitative susceptibility reconstruction algorithms.

## 1 Introduction

Quantitative Susceptibility Mapping (QSM) has proven to be a valuable tool for assessing iron concentrations in the deep gray matter (DGM) (1–3), estimating vessel oxygenation and geometry (4,5), differentiating blood and calcium products (6,7), and studying demyelinating lesions in the white matter (WM) (8–11). However, several recent methodical investigations have suggested that study outcomes may depend on the particular processing algorithms chosen for QSM (12–14). QSM typically involves the following steps in the order of application: coil combination (12); phase unwrapping (14); multi-echo combination (12); background field removal (14); and, finally, the estimation of susceptibility maps (13,15–17). Processing artifacts and inaccuracies at any of these five processing stages can propagate into the computed susceptibility maps.

The first QSM Reconstruction Challenge (RC1) in 2016 (18) aimed to provide initial insights on the accuracy of various proposed algorithms for estimating susceptibility from background-corrected frequency maps, i.e., the last processing step of QSM. One of the key conclusions of RC1 was that the choice of the algorithm and the used parameter settings can have a substantial, non-negligible effect on the appearance and accuracy of computed susceptibility maps. However, upon completion of the challenge, it was also recognized that the particular gold-standard (reference) susceptibility maps used for evaluating the challenge submissions limited the interpretability of the challenge outcomes. The reference maps were generated from multiple acquisitions in which the subject had rotated the head towards 12 different orientations. From these data, two reference maps were created, one calculated with the susceptibility tensor imaging (STI) (19) technique and one by Calculation Of Susceptibility through Multiple Orientation Sampling (COSMOS) (20). Meanwhile, only one of the 12 field-maps was provided to the challenge participants. The rationale of this approach was that RC1 would yield the most objective and meaningful results if algorithms were evaluated using real-world in vivo data. However, at the completion of RC1, it was observed (21) that a non-negligible discrepancy existed between the provided frequency map and the frequency map obtained when the field perturbation was forward-simulated based on the provided reference susceptibility maps. It was speculated that a part of the discrepancies were related to unaccounted microstructure effects on in vivo brain phase images (22). Current single-orientation QSM algorithms assume that frequency contrast is caused entirely by variations in bulk magnetic susceptibility and all other contrast mechanisms are neglected. Consequently, the discrepancy between the provided field map and the gold standard susceptibility reference rendered it challenging or even impossible to achieve a reconstruction from the field map that was close to the reference used. It turned out that the best performing RC1 submissions, i.e. those with the smallest error metrics, were over-regularized and had a non-natural appearance.

The goal of the second reconstruction challenge (RC2) in 2019 was to address the identified limitations of RC1 and provide more meaningful insights on the current state-of-the-art in QSM algorithms to identify their strengths and limitations in different scenarios and inform and coordinate future methodological research efforts. During the planning phase for RC2, the challenge committee concluded that the systematic evaluation of the accuracy and robustness of QSM methods should focus on synthetic (in silico) phantoms with realistic forward simulations rather than on real-world data. The challenge was designed with two stages: Stage 1 mimicked the clinical setting in which the ground-truth was unknown to participants; in Stage 2 the ground-truth was made available and, thus, allowed for systematic parameter optimisations to obtain the *best possible* quality metrics that can be obtained with each reconstruction algorithm. The results of RC2 were reported in a separate manuscript (23).

In this paper, we present the modular framework designed to generate the realistic digital head phantoms for RC2. Methodological researchers may use the RC2 phantom in their studies to evaluate existing and future QSM algorithms and compare their results to RC2 submission. The comparison of their metrics to those of RC2 submissions will facilitate the objective evaluation of methodological improvements and algorithm performance across labs. As more advanced physical models are incorporated into the QSM algorithms, researchers may extend the phantom according to their needs. Code and data are freely available and have been designed to facilitate adding new features to the phantom, such as calcifications and hemorrhages or microstructure effects. The software package may also be used to optimize acquisition protocols, prepare and test complete QSM reconstruction pipelines. In combination with other software, the package will allow evaluating the effect of image distortions or blurring on QSM.

## 2 Methods

### 2.1 Design considerations

#### 2.1.1 Limitations of previous evaluation strategies

In the literature, most QSM algorithms were evaluated based on their visual appearance (5,24,25), based on the root-mean-squared-error (RMSE) of reconstructions of simple digital piece-wise constant phantoms consisting of geometrical shapes (25–30), or simplistic head phantoms (22,31). Evaluation of the susceptibility quantification accuracy and precision typically relied on phantoms made of agar or aqueous solutions with varying concentrations of contrast agents such as Gadolinium (20,26,32–35) or iron oxide particles (5,28,36–38). Such measurements have been of great importance to establish that QSM linearly maps the magnetic susceptibility property and that measurements across different platforms can be compared. In vivo, QSM accuracy has often been evaluated (27,31) by using previously published iron concentrations in the deep grey matter nuclei (39) as a surrogate gold standard. This approach suffers from the large variability in iron concentrations across subjects.

A major limitation of most previously used digital phantoms and liquid or gel phantoms was that they had piece-wise constant susceptibility distributions, which are particularly easy to invert for methods with total variation regularization (“inverse crime”). To address this limitation, validation has also been performed by injecting gadolinium into tissue samples (20) or using air bubbles or glass beads (40). In the first case experiments, however, the ground truth is again not available as the agents diffuse within the tissue. Therefore, it has to be reverted to visual inspection. In vivo, as an alternative to visual inspection, maps have been compared to a COSMOS reconstruction of the same subject (28–30,41,42) due to their reduced level of streaking artifacts, similar as in RC1. However, using a COSMOS solution as gold standard implicitly assumes that the measured phase satisfies COSMOS’s field model. Specifically, COSMOS assumes that (i) susceptibility is isotropic throughout the brain; (ii) that the dipole model with the sphere of Lorentz approximation can be used throughout the brain (22); and (iii) that microstructure-related frequency effects (43) and chemical exchange (44,45) do not exist. Since these assumptions are simplistic, COSMOS does not generate an appropriate ground truth susceptibility map for single-orientation QSM and, hence, cannot be considered a good gold standard method.

#### 2.1.2 Design considerations for RC2

This RC2 committee’s decision to employ a digital phantom resulted from the realization that a true gold standard technique for in vivo QSM did not exist. Without a gold standard technique, it was not possible to obtain a meaningful in vivo reference susceptibility map through measurements. The committee had also discussed the design of *real-world* test objects (phantoms) that are consistent with the QSM phase model (33,34). Based on the committee members’ experience with phantom design and a literature research of previously used phantoms, it was concluded that the inclusion of sufficiently complex morphology and fine-scale susceptibility features would be prohibitively challenging. It was unanimously concluded that it would be most reasonable to focus the committee’s efforts on a digital phantom that could be adapted and extended to the community’s evolving needs in the future. For real-world data, available references usually represent only approximations of the ground truth (gold standard). On the contrary, *in silico* phantoms provide a genuine *ground truth.* In silico phantoms also allow for a controlled investigation of the effect of deviations from the underlying QSM model on the reconstruction performance. Besides the ability to model different biophysical phase contributions, digital models also allow a controlled inclusion of measurement-related phase errors. For example, field measurements close to the brain surface are affected by nuisances, such as signal drop-out and the non-linearity of the phase evolution due to the non-negligible higher order spatial terms inside the pixel (46) that make the measured field deviate from the actual voxel-average field. Similar limitations are present when developing background field removal methods. Despite this known limitation, only a few methods have the possibility to explicitly account for field map uncertainty (27,32), while remaining methods address this problem by increasing the brain mask erosion (24,26,47).

As a first step toward a future systematic evaluation of all above-mentioned experimental aspects influencing QSM reconstruction quality, the RC2 *in-silico* phantoms enforced consistency of the provided frequency map with the physical model used by current QSM algorithms. Also, the RC2 phantoms were designed to feature a realistic brain morphology and naturally varying susceptibility distribution within anatomical regions.

### 2.2 Data acquisition

We acquired MRI data from a human volunteer (F, 38 years) who gave informed consent, and the experiment was approved by the local Medical Ethical Committees (Amsterdam University Medical Centre and Radboud University Medical Centre). We used a 7T scanner to obtain relaxation rate maps and a 3T scanner to obtain DTI data and bone-air tissue interfaces. For generating the brain phantom, we acquired inherently co-registered quantitative maps of R_1_ (48), R_2_*, χ and M_0_ maps using the MP2RAGEME (49) sequence on a 7T (Philips Achieva) scanner. The main sequence parameters were TR/TI_1_/TI_2_=6.72/0.67/3.86 s. The first and second inversion times were acquired with TE_1_=3ms and TE_1/2/3/4_=3/11.5/20/28.5ms and flip angles α_1_/α_2_ = 7/6 °, respectively. The acquisition was performed sagittally with FOV=205×205×164 mm^3^ and matrix size = 320×320×256, resulting in an isotropic resolution of 0.64mm and its total acquisition time was of 16min 30secs.

For generating a bone and air model, we acquired T_1_w (0.93 mm isotropic) data at very short echo time using the PETRA (50) sequence with the following parameters at 3T (Siemens PrismaFIT): TR_1_/TR_2_/TI_1_/TI_2_=3/2250/1300/900 ms; flip angle=6°, TE=0.07 ms; matrix size = 320×320×320; total acquisition time 5min 57secs.

To add microstructure effects to the phantom, we acquired DTI data using two simultaneous multi-slice EPI-based datasets with opposed phase encoding directions. The main sequence parameters were TR/TE=3520/74 ms, SMS factor 3, in-plane acceleration 2, matrix size = 140×140×93 and FOV=210×210×139.5 mm^3^, resulting in 1.5mm isotropic image resolution. The diffusion-weighted parameters were b=0/1250/2500 s/mm² and 12/90/90 directions, respectively, resulting in a total acquisition time of 12 min 10 secs. Diffusion data was processed using fsl software (https://fsl.fmrib.ox.ac.uk/fsl/fslwiki/), eddy_correct and top up were used to undistort the DWI data. Data was coregistered to the 7T anatomical space and FMRIB Diffusion Toolbox was used to extract tensor information (FA, fractional anisotropy, main eigenvector orientation).

### 2.3 Tissue segmentation

Figure 1 shows a pictorial representation of the pipeline used for the segmentation. The 7T T_1_ maps, derived from the MP2RAGE dataset, were segmented into 28 tissue classes, including left and right splitting, using the cbstools atlas based pipeline (https://www.cbs.mpg.de/institute/software/cbs-tools(51)). Classes were then re-clustered into 16 tissue clusters: CSF (initially split into 4 classes); grey matter (initially split into 8 classes, left and right cortical, cerebellar, amygdala and hippocampus); caudate; putamen; thalamus; white matter (encephalus, cerebellum and brain stem); and large blood vessels. Deep grey matter structures not clearly distinguishable on T_1_ maps (red nucleus, substantia nigra, globus pallidus and dentate nucleus) were manually segmented using the active contours function implemented in ITKSnap (version 3.6) (52) on R_2_* and χ maps. A calcification present in the subject’s interhemispheric fissure was identified and segmented using the M_0_ map. An initial vein-and-artery mask was computed based on Frangi filtered R_2_* maps as described in (53). The region outside the brain was segmented into bone, air and tissue using a model-based segmentation approach with deformable surface meshes (54) using the PETRA sequence as input. The resulting segmentations of nasal cavities and auditory canals were refined manually using ITKSnap for the computation of realistic background fields (see below).

**Figure 1.**
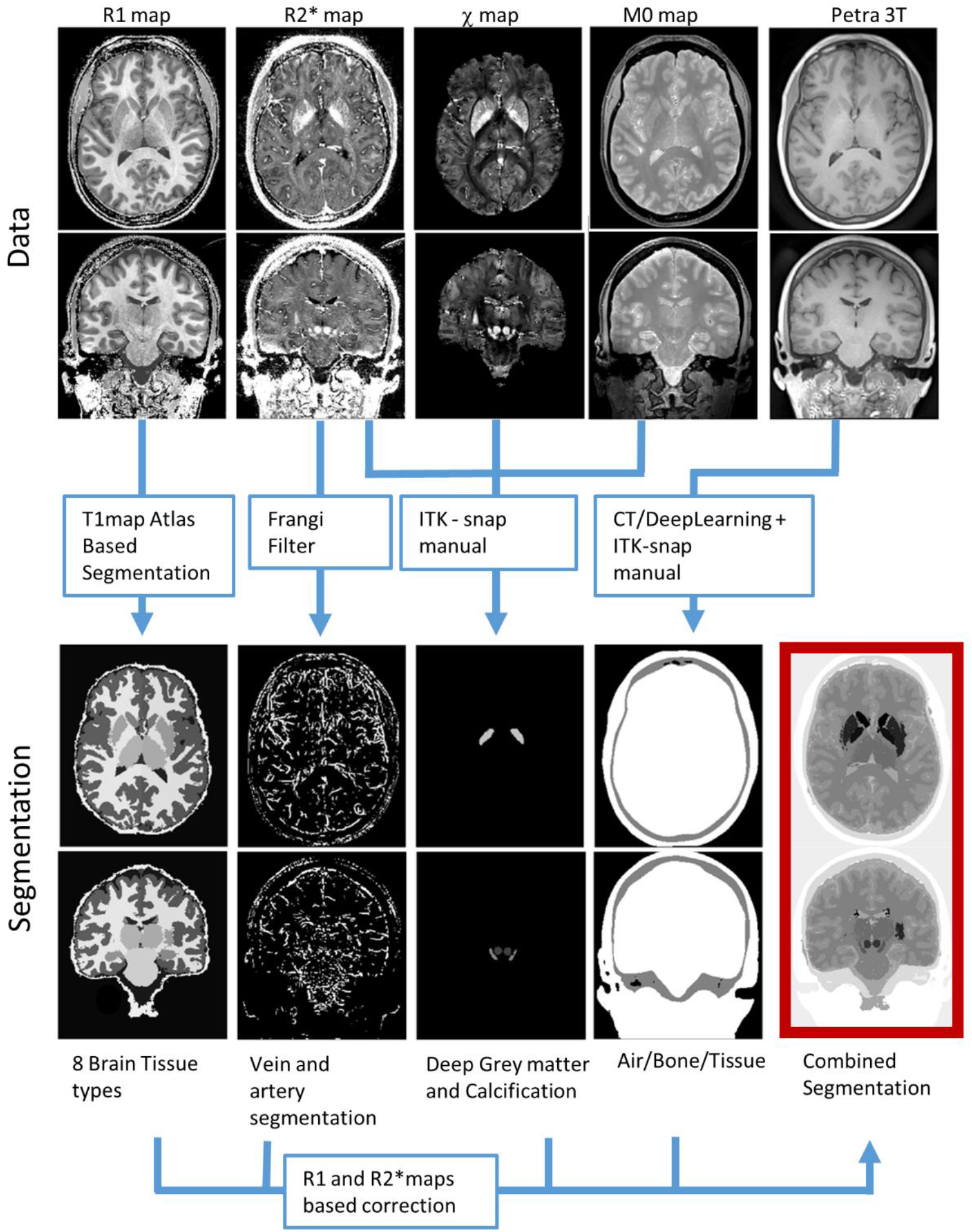
Diagram representing the process used to obtain the head segmentation: R_1_ map (obtained from the MP2RAGEME) was used to create an atlas-based segmentation using CBS tools; R_2_* were processed with a Frangi filter for vein segmentation; A semi-manual approach using ITK snap was used for segmentation of the deep gray matter nuclei based on the R_2_* and a susceptibility map computed using HEIDI, and the M0 map were used to segment the calcification. Finally, PETRA data was used to obtain air, bone, and tissue masks using a CT-based deep learning algorithm followed by manual ITK-snap. Then, the various tissue segmentations were fine-tuned using denoised R_1_ and R_2_* maps using manually defined thresholds. The various masks were combined to generate a whole-head segmentation with 16 different tissue types.

After the combination of the individual tissue brain masks into a piece-wise constant whole-head phantom, we used R_2_* and R_1_ maps to correct the label boundaries in various brain regions using customized thresholds (see Table S1 in the Supplementary Material). The initial tissue segmentations were based on one single quantitative parameter (R_1_ for tissues compartments, R_2_* for veins), and because smooth surfaces had been enforced in some regions this resulted in segmentation mismatches that benefited from this second processing iteration.

### 2.4 Susceptibility map

The susceptibility map was simulated by assigning tissue-typical susceptibility values taken from literature (55,56), 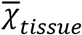 (see Table 1), to the various tissue segments. We modulated the susceptibility values in each region using the image intensities on R_1_ and R_2_* maps according to the following equation:

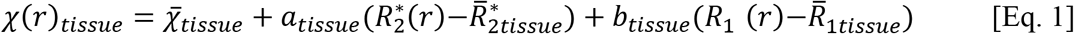

where 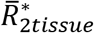 and 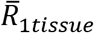 are the mean apparent transverse and longitudinal magnetization rates of that given tissue segment class. There were three main motivations for using such an expression to compute our ground truth susceptibility map:

- Modulation avoided that susceptibility values were constant throughout anatomical regions (piece-wise constant). Absence of modulation would be both unrealistic and advantageous to algorithms with gradient-based regularization terms.
- From a practical perspective, R_1_ and R_2_* were the only two “bias field” free maps available at high resolution that could be used to create a anatomically valid intensity modulation;
- Both transverse and longitudinal relaxation rates, like magnetic susceptibility, are known to have a linear dependence on the concentration of paramagnetic and diamagnetic perturbers when dealing with simple liquid solutions. The main difference to susceptibility is that relaxation rates are agnostic to the sign of the magnetic perturber; particularly in brain tissues, both R_1_ and R_2_* have been shown to have a linear dependence on the concentrations of iron and myelin (57).

**Table 1.**
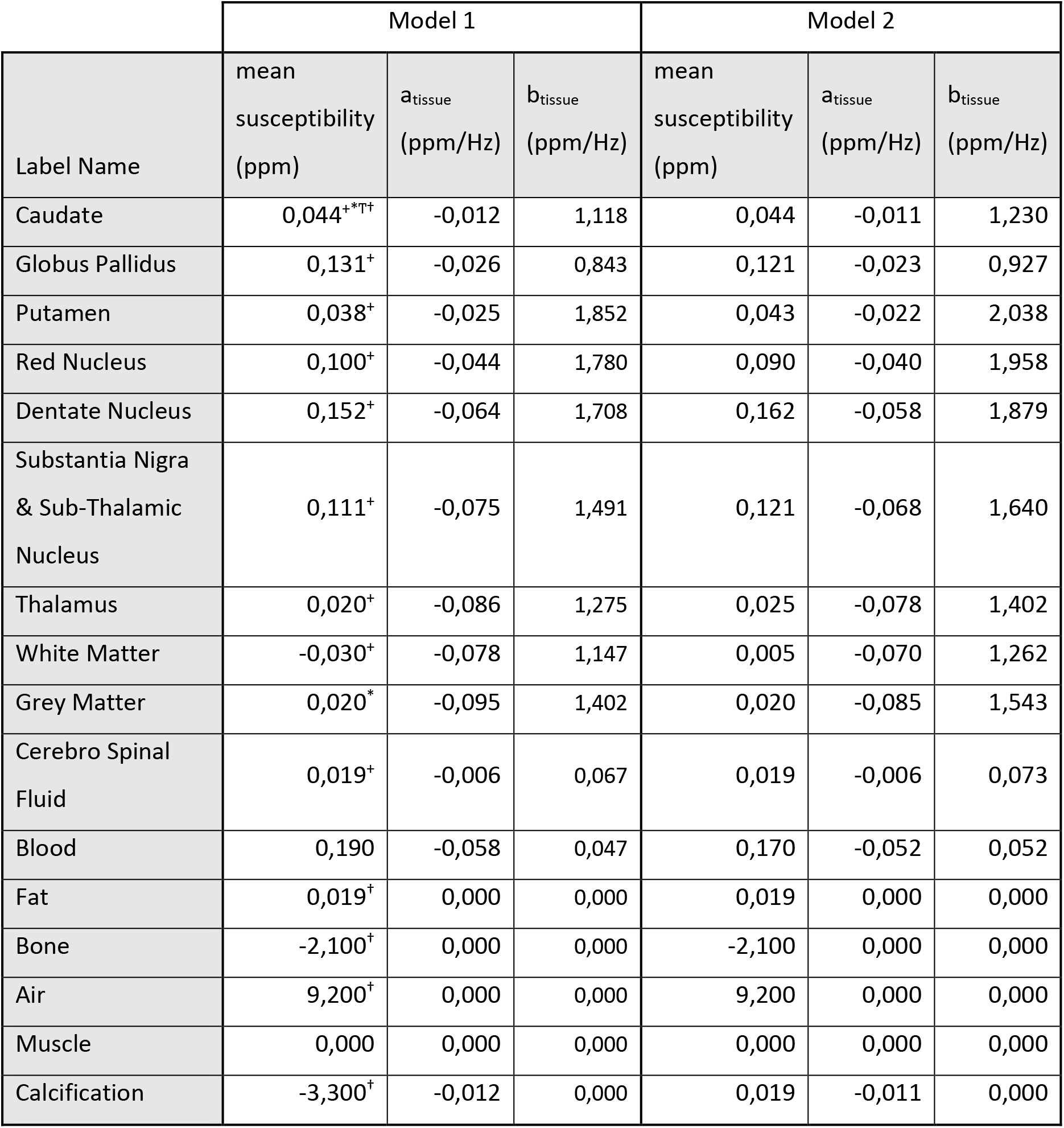
Parameters used to create the two magnetic susceptibility head models released in the QSM challenge. The values in the three columns correspond to the parameters described in equation 1, the assumed mean magnetic susceptibility of the tissue, and the R_2_* (a_tissue_) and R_1_ (a_tissue_) modulation terms, respectively. Model 1 values were chosen from literature (^+^ (55), ^*^ (22) ^†^ (56)) and doing the fitting described in Eq. 1. Model 2 mean susceptibility values were adhoc modification of those found in literature, while the R_1_ and R_2_* modulation were changed by plus and minus 10% respectively.

While it is reasonable to assume that the susceptibility map (assuming any other tissue properties constant) could be given by a linear combination of these two maps, the main aim was to introduce a realistic texture. To obtain proportionality parameters, *a*_*tissue*_ and *b*_*tissue*_, resulting in realistic susceptibility variations, Eq.1 was inverted for each brain tissue class using as χ(*r*)_*tissue*_ the HEIDI susceptibility map calculated from the original data. The coefficients 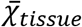, *a*_*tissue*_ and *b*_*tissue*_ used for the two phantoms in the QSM challenge 2.0 can be seen in Table 1 and Figure 2. Note that having different proportionality parameters for each tissue (in addition to a different mean value per tissue type), results in a susceptibility map that cannot be derived simply from the magnitude signal variations. Because the measured relaxation rates of blood (both in arteries and veins) are prone to errors (due to inflow effects on R_1_ and flow effects on R_2_* maps), the proportionality values were relatively small for the blood pool, rendering these compartments piece-wise constant. Bone, calcification and other non-brain tissue compartments were made piece-wise constant (the lack of a susceptibility map outside the brain prencented the derivations of *a*_*tissue*_ and *b*_*tissue*_). Close to air-tissue boundaries, where strong field gradients are present, tissue R_2_* values are overestimated (see R_2_* maps in Fig.1 over the ear canals) (58). A low-pass filtered version of the gradient of the acquired field map was used to differentiate regions where the R_2_* values could be trusted from those where they were unreliable. In the latter regions, we forced *a*_*tissue*_ = 0 (ignore R_2_* contribution when generating the susceptibility phantom). To avoid discontinuities between high (>0.08ppm/mm) and low (<0.3ppm/mm) field gradient regions, a smooth transition was created by mixing the two combinations (from full Eq.1 and *a*_*tissue*_ =0, respectively). Please refer to provided code for more details on the implementation.

**Figure 2.**
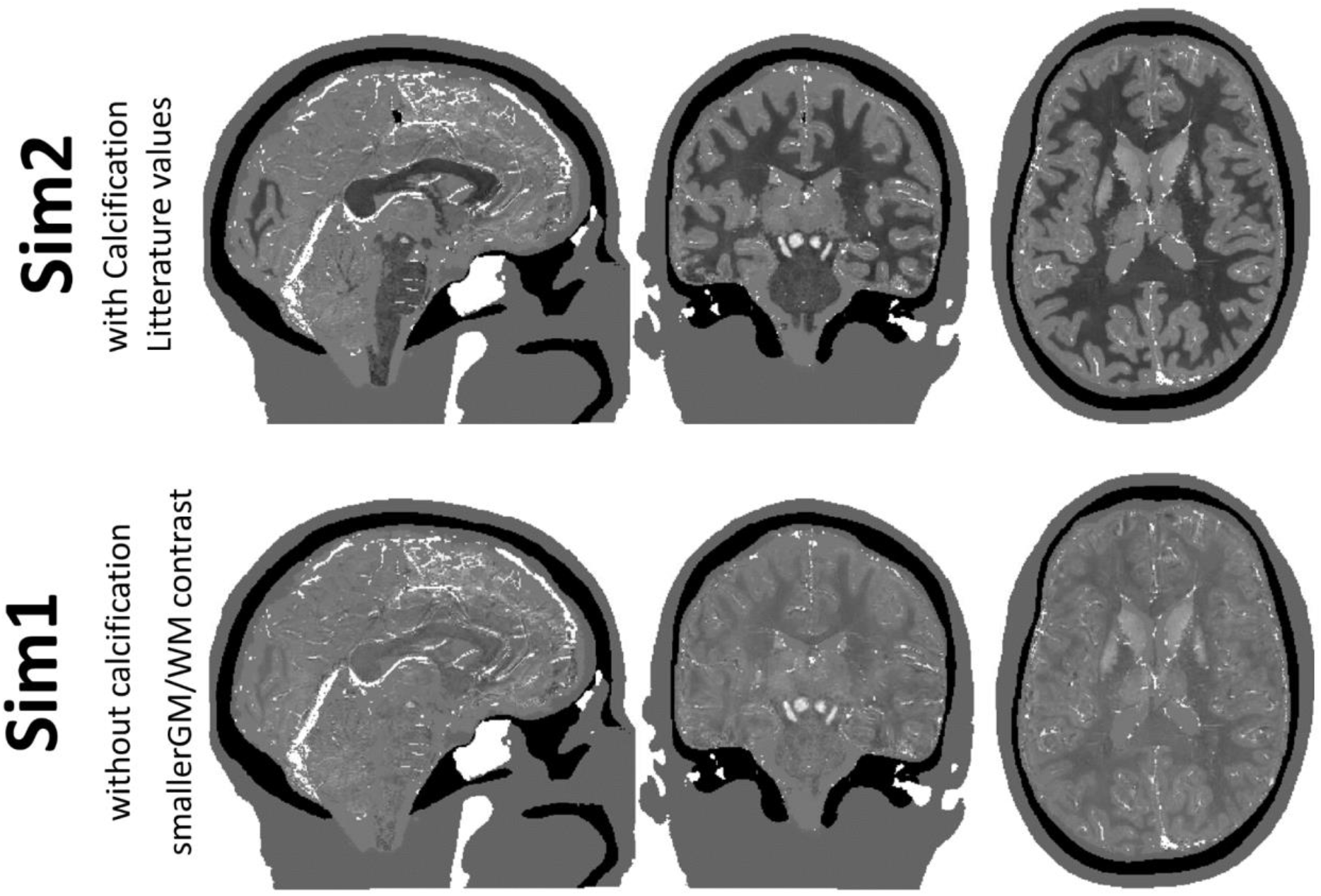
View of three slices in sagittal, coronal and axial directions of the two digital phantoms created for the QSM challenge 2.0. Top and Bottom maps were obtained using the parameters described on Table 1 for Model 1 and 2 respectively.

To avoid unrealistically sharp edges of magnetic susceptibility at the interfaces between tissue regions, we have introduced partial voluming in those transitions. Note that transitions between brain tissues tend not to be sharp (for example, grey matter layers on the white matter side are highly myelinated (59), as are the outer parts of the thalamus), while between tissues and blood, CSF, air, bone and muscle the interfaces will be sharp. The probability of a voxel being a given tissue, P_tissue_, was computed by smoothing each binary brain tissue mask using a 3D Gaussian kernel with a full-width at half maximum (FWHM) of 1.2 voxels. This smoothing was not applied to veins or non-brain tissue masks. The probability was computed as 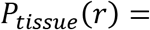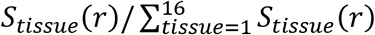, where *S*_*tissue*_ is either the smoothed or unsmoothed mask of a given compartment depending on it being a brain tissue or non-brain tissue. Note that transitions between brain tissues tend not to be sharp (for example, grey matter layers on the white matter side are highly myelinated, as are the outer parts of the thalamus), while between tissues and blood, CSF, air, bone and muscle they will be sharp. The susceptibility phantom was then given by:

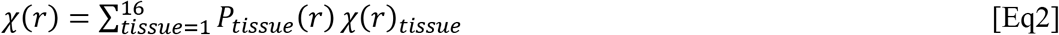

The importance of moving from a piece-wise constant (where a *a*_*tissue*_ and *b*_*tissue*_ are set to zero) to a contrast-modulated (where *P*_*tissue*_ is simply a binary mask) or the probabilistic formalism of the susceptibility distribution can be appreciated in Figure 3. Yellow arrows highlight the transition between grey and white matter that becomes smoother, and green arrows smoothing out small segmentation errors within the thalamus. On the other hand, Figure 3 shows that most vessel structures have remained sharp with only some minor reduction in susceptibility value. R_2_* maps tend to enlarge venous structures due to blooming artifacts. By not smoothing the blood compartment mask when computing the final susceptibility map, this effect was not further extended. On the other hand, because neighboring tissues have been smoothed into the blood compartment, the blood partial volume in blood vessels is reduced resulting in a lower susceptibility the smaller the vessel is, mimicking a realistic scenario.

**Figure 3.**
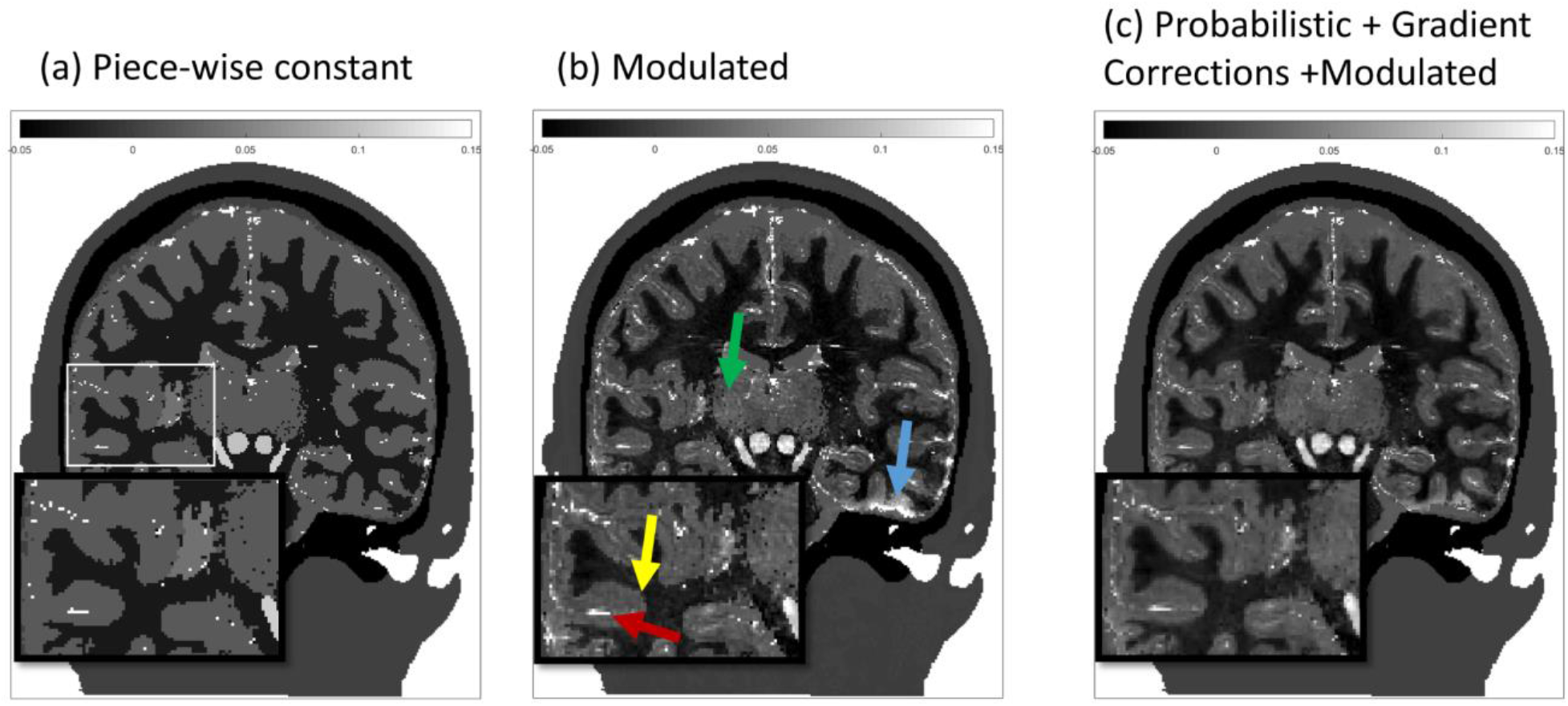
Intermediate stages of the of the creation of the in silico susceptibility head phantom: (a) traditional piece-wise constant approach, (b) modulated model as described in Eq. 1 with the values presented in Table 1 and (c) finally adding the probabilistic modulation described in Eq. 2 and masking regions of error bound R2*. Green and yellow arrows highlight transitions between tissue types that get improved via the probabilistic approach applied to the tissue compartments. Note that the probabilistic smoothing was only applied to the brain tissues, as a result, veins remain shape in the susceptibility map only with a reduced magnetic susceptibility. Blue arrow highlights regions where the large field gradient masking approach was able to avoid abnormally large susceptibility values.

### 2.5 Data Simulation

Spoiled gradient recalled echo data can be simulated using the steady state equation:

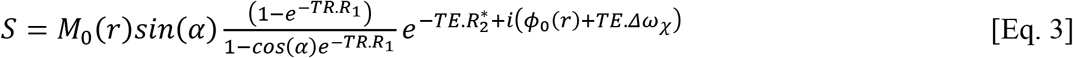

Here, ϕ_0_ is an initial phase distribution originated from the transceiver phase, and *Δω*_χ_ is the frequency shift directly attributed to magnetic susceptibility. To simulate S for given repetition times (TR), echo times (TE), and flip angles (*α*), we used the M_0_, R_1_ and R_2_* maps derived from the MP2RAGEME (49,60) sequence, ϕ_0_(*r*) is the TE=0 phase (a manually selected 3D second order polynomial ensuring 2π phase variation inside the brain region was used), and the frequency shift, Δω(r), was calculated according to

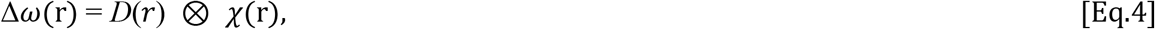

where *D*(r) is the magnetic field dipole along the z-direction with Lorentzian correction. The convolution was performed in k-space using the formulation proposed in (61,62). To avoid aliasing artifacts associated with the Discrete Fourier Transform (circular convolution), which would appear as unrealistic background fields, we padded the phantom with zeros along each dimension (factor of 2) before evaluating Eq. 4. Such a formulation of the signal equation, explicitly neglects chemical shift (associated with spins from, for example, fat) and any chemical exchange effects on the frequency. Furthermore, because no gradient waveform was defined explicitly, image distortion effects were not simulated, and the impact of blood flow on phase data was not accounted for.

A digital phantom enables simulating the MR signal with and without background fields (fields generated by tissues and other sources located outside the brain). The latter effectively mimics “perfect” background-field correction. Using the whole-head susceptibility phantom allowed the creation of realistic background fields. To create a phantom without background fields, named hereafter “local field”, all voxels outside the brain were set to zero and the susceptibility distribution within the brain was demeaned:

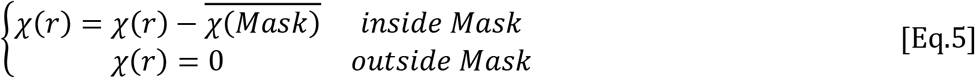

Also relevant, but not pursued for the QSM challenge purposes, field shimming was simulated by fitting the frequency map with second and third-order Legendre polynomials.

#### 2.5.1 Simulation of different acquisition protocols

For the demonstration of the acquisition protocol simulation with the code described in supplemental material, we chose two example protocols designed for different applications:

(P1) A protocol optimal for the quantification of deep grey matter susceptibility. In this case, the longest TE should be at least that of the T_2_* of the region with the highest iron concentration which was the globus pallidum in the generated phantom (14ms).
(P2) A protocol optimal for the observation of cortical grey/white matter contrast in both the magnitude and phase data. In this case, the longest TE was chosen to be close to that of the T_2_* of cortical GM (33ms) (63).

For the sake of simplicity, both protocols had the same echo spacing of 8ms as the original volunteer dataset. The TR of the acquisition was chosen as short as possible assuming a readout acquisition window of 8ms (P1: TR = 16ms; P2: TR = 40ms). We neglected dead times associated with phase encoding preparation, flow compensation, rewinding, rf excitation, saturation, and crushers. The flip angle was chosen at the Ernst angle for the GP (T_1_=1100ms; α=8) and such that T_1_-weighted contrast on magnitude images was maximized between WM (T_1_=1100ms) and cortical GM (T_1_=1900ms; α=23) in protocols P1 and P2, respectively. This resulted in the following protocols:

Protocol 1: TE_1_/TE_2_= 4/12ms;
Protocol 2: TE_1_/TE_5_= 4/36ms.

We mimicked k-space sampling by cropping the Fourier spectrum of the original 0.65mm resolution data to an effective spatial resolution of 1mm isotropic. We applied the same approach to down-sample the ground-truth susceptibility map. In the case of the susceptibility maps, the sharp edges between structures as well as the orders of magnitude larger susceptibility differences between air/bone and tissue resulted in severe Gibbs ringing artefacts, which were removed using sub-voxel shifts (64). This step was repeated in all three spatial directions.

Further processing consisted of spatial unwrapping of echo differences using SEGUE (65), combination of resulting field maps using the optimum weights (12,41), and (when necessary) removal of background fields using the Laplacian Boundary Value (LBV) method (26).

#### 2.5.1 QSM reconstruction optimization

To demonstrate the applicability of the current framework for the QSM challenge or for QSM reconstruction optimization purposes, we performed a simulation with only local fields (see Eq. 5). The protocol used was that of the QSM challenge (TR = 50ms; TE_1/2/3/4_= 4/12/20/28ms; α=15, (23)). QSM reconstructions using TKD (66), closed form L2 (67), FANSI (68) and iLSQR (69) as implemented in the SEPIA toolbox (70) were performed with varying regularization parameters. The reconstructions were evaluated using the reconstructions metrics created for the challenge (see Table 2). For a more detailed description refer to Supporting Information (Section 2).

**Table 2.** Metrics provided with toolbox and challenge for optimization and evaluation of QSM reconstruction. Note that all the RMSE metrics were multiplied by 100

| Metric Name | Description |
| --- | --- |
| nRMSE | Whole-brain root-mean-squared error after demeaning, i.e. the subtraction of the mean within the mask; relative to the ground-truth times 100; |
| rmse_detrend_Tissue | nRMSE relative to ground-truth (after demeaning and detrending) in grey and white matter mask. Detrending was performed by compensating the estimated the slope of a linear fit of the reconstructed QSM voxel values against those of the ground truth in the tissue ROI. The reconstruction was then divided by this factor to ensure proportionality errors (measured by other metrics such as DeviationFromLinearSlope) do not affect the RMSE calculation; |
| rmse_detrend_blood | RMSE relative to ground-truth (after detrending) using a one pixel dilated vein mask; |
| rmse_detrend_DGM | RMSE relative to ground-truth (after detrending) in a deep gray matter mask (substantia nigra & sub-thalamic nucleus, red nucleus, dentate nucleus, putamen, globus pallidus and caudate); |
| DeviationFromLinearSlope | Absolute difference between the slope of the average value of the 6 deep gray matter regions vs the prescribed mean value and 1.0. |
| CalcStreak | Estimation of the impact of the streaking artifact in a region of interest surrounding the calcification by means of the standard deviation of the difference map between reconstruction and the ground-truth. The region of interest where this was measured was a hollow rectangular prism, with its inner boundary being 2 voxels away from the edge of the calcification and the outer boundary 6 voxels away from the inner boundary. |
| CalcMoment | volumetric susceptibility moment of the reconstructed calcification, compared to the ground-truth (computed in the high resolution phantom to be -49.8 ppm). This metric has been suggested to be more robust in regions of punctuated large susceptibility sources where there is no signal in the region of the perturber (40). |

#### 2.5.2 Adding microstructural effects to the obtained contrast

Microstructural effects are known to affect the observed phase. One of the driving factors of the microstructural effects is white matter fiber orientation. The provided data and code include a simple first-order approximation of these microstructure effects which is echo-time independent. Wharton et al (22) demonstrated that the typical impact at 7T for a protocol with echo times of 7 and 13 ms was given by:

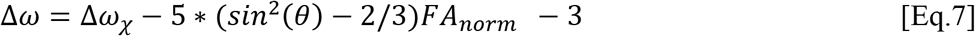

where FA_norm_ is the fractional anisotropy (FA) divided by 0.59 (the average anisotropy observed in a human optic nerve) and θ is the angle between the white matter fiber and the static magnetic field. Both of these quantities can be derived from the acquired diffusion data. Such a correction to the frequency shift was only applied within the segmented white matter mask.

## 3 Results

### 3.1 Simulations of different acquisition protocols

Figure 4 shows Phantom 1 with the two different sequence parameters created by the proposed simulation toolbox (See Supporting Information Section 1 or data sharing collection for code). The top row shows an example slice of the simulated data from the P2 protocol aimed at computing QSM in cortical grey and white matter. In this case, the longer echo time matched the T_2_* of those grey matter resulting in both significant signal decay in deep gray matter and a large number of phase wraps close to tissue air boundaries. The flip angle used (23 degrees) was set to increase T_1_ contrast in grey vs white matter boundaries at short echo times as can be clearly appreciated on the top left figure. Such information can be used to inform the QSM algorithm regarding expected morphological features. The bottom row shows the images associated with the P2 protocol, aimed at measuring the susceptibility values in deep grey matter regions. The echo time range is smaller, so that the magnitude signal in the globus pallidus has not yet disappeared at the last echo time and the tissue contrast in the magnitude data is considerably weaker because of the smaller flip angle (8 degrees). From the data shown in Figure 4 it can be expected that P1 will benefit more from a morphological informed reconstruction than P2, and that the use of non-linear fit (71) for the calculation of the field map will also be particularly relevant as noise will dominate the later echoes in P1.

**Figure 4.**
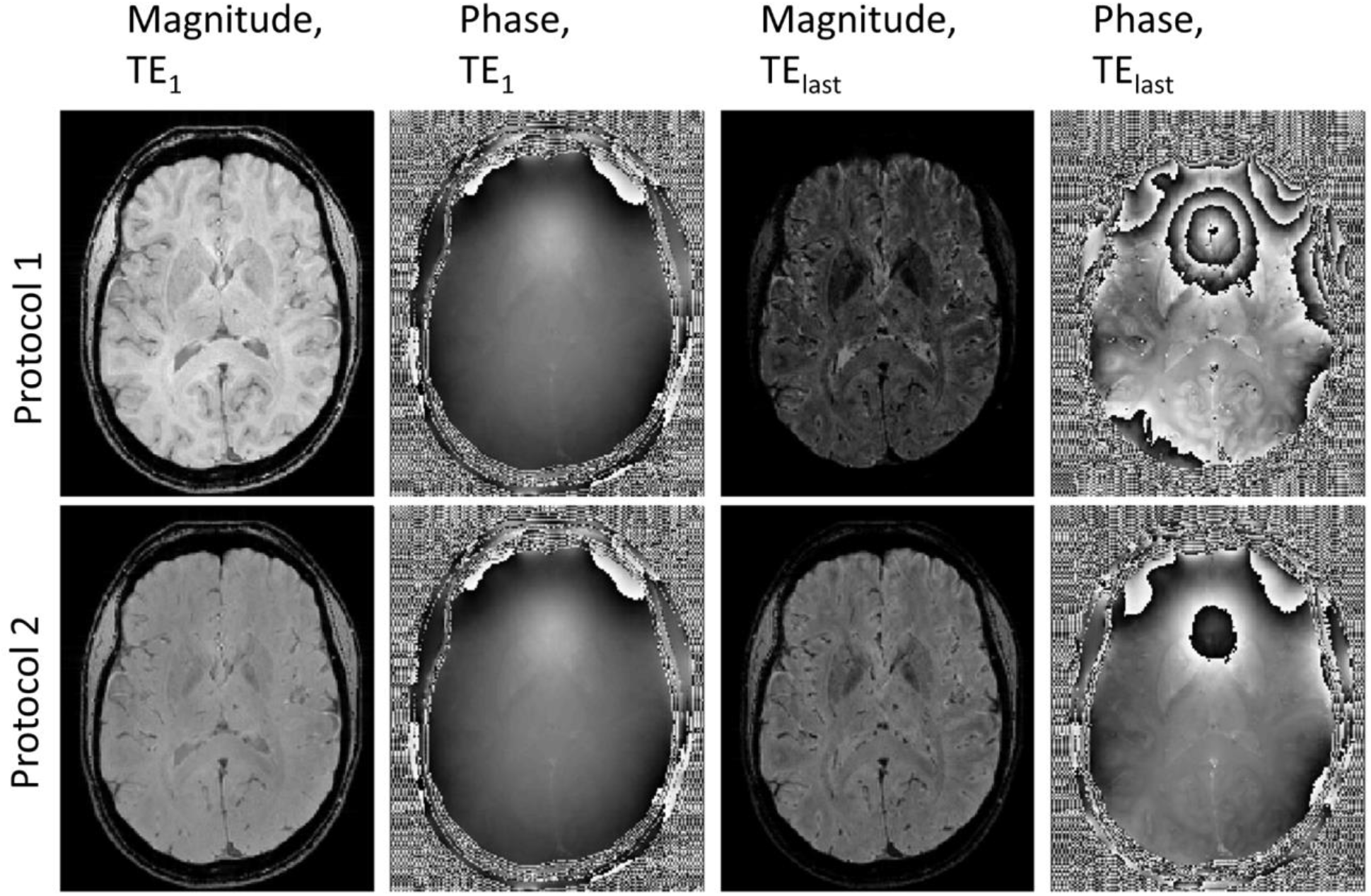
Transverse slices of the simulated data of Model 1 using Protocol 1 (top row) and Protocol 2 (bottoms row). First two columns show magnitude and phase images, respectively, at the first echo time (where the different T_1_-weighting is clearly visible) and the last two columns show the magnitude and phase images associated with the last echo time of the respective protocol.

In Figure 5 we compared the field map obtained with the P1 simulation from the whole head phantom to a field map obtained directly from the susceptibility brain phantom via Eq. 5 (ground truth field map). When carrying out the signal simulation with the whole-head phantom, both the base resolution and the 1mm resolution field maps are dominated by the background components arising from the air/bone/tissue interfaces, as can be seen in the transverse slice above the sphenoid sinus (first column, as pointed by the black arrow). The differences to the ground-truth field map (second column) are to a large extent explained by the quadratic field used to represent the procedure of shimming. Once LBV was applied to the total field to obtain the tissue-specific field contributions (third column), the original resolution field map (top row) showed localized, smoothly varying differences relative to the ground truth, which have been described previously (72). Both the base resolution and 1mm resolution look at first sight to have very similar properties, yet the nRMSE from the down-sampled dataset (bottom row) demonstrated additional deviations from the ground truth (nRMSE was 40% higher than that of the high-resolution dataset). This discrepancies are both due to incomplete background field removal (see grey arrows) and due to errors around veins. In such regions, the field map measured from the GRE data naturally deviates from the mean field in that pixel because of the reduced signal in veins (once partial volume is introduced by the reduced resolution, the field estimation is biased towards the tissue compartments).

**Figure 5.**
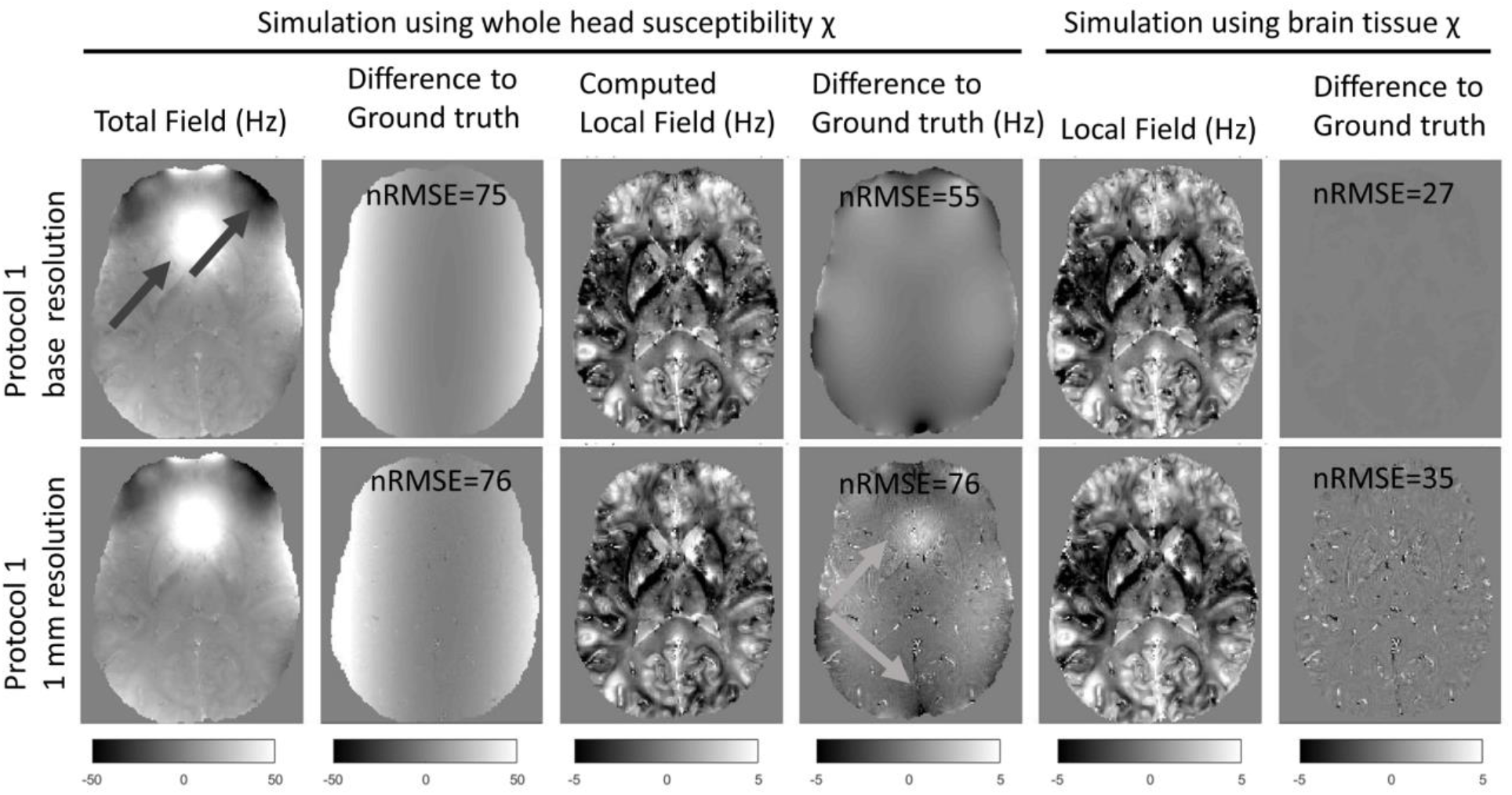
Transverse slices through derived field-maps associated with Protocol 1 data computed at the base resolution (top row) and after down-sampling to 1mm (bottom row), black arrows highlight large background fields induced by air-tissue interfaces. The first and fifth columns shows the field extracted from the complex signal when the whole-head and the brain-only models were used, respectively, to compute the frequency shift. The third column shows the local field computed after background field removal. The second, fourth and sixth columns show the differences relative to the corresponding ground-truth field distributions. Grey arrows highlight the incomplete background field removal that is exacerbated upon downsampling. The ground-truth field maps were computed using the forward dipole formulation (Eq. 4) on the whole head (second column) and brain tissues alone (fourth and sixth columns) susceptibility models. It is clear that the nRMSE, once the down-sampling is performed, is dominated by partial volume effects in and around veins (noise pattern on both bottom 4^th^ and 6^th^ column) and imperfect background field removal.

To disentangle the effects of background field correction and MRI signal simulation on the deviations observed relative to the ground truth, we repeated the signal simulation with the local fields. In that case, the slowly varying smooth deviations disappeared and the high-resolution phantom did not demonstrate substantial deviations relative to the ground truth. The differences in the field computed at base resolution without background fields (top right panel) is at the numerical precision level, yet the nRMSE is still not negligible (nRMSE = 27) because of the errors present in the calcification region without signal exists and its immediate surrounding where spatial unwrapping fails.

To further investigate the sources of errors discussed in the previous paragraph, Figure 6 evaluates the phase evolution in three voxels, two in the surrounding of the calcification and one in the white matter. Figure 6c shows in the case of an homogeneous tissue region, there is a perfect match between low resolution and high resolution phase evolutions as well as the fitted frequency (based on the 5 echo simulation) and the ground truth frequency (computed from the susceptibility map). Figure 6b shows that in a region closer to the calcification that is also the case for the high-resolution data (light grey lines), while in the case of the low-resolution data (dark grey) there is a larger error both in respected to fitted frequency (dashed line) and ground truth frequency evolution. Also, it is clear that the phase evolution in Fig. 6b in the low resolution data is no longer linear, as is predicted from theory due to partial volume effects and the varying intravoxel frequency gradients (46). In a pixel immediately close to the vicinity of the calcification the errors are further enhanced now also for the high resolution data where unwrapping errors can introduce errors on the fitted frequency (the data is still fitted accordingly, but it does not correspond to the ground truth field). Figure 6e shows the mean squared difference map associated with the frequency fit on the low resolution data, it is clearly visible that these errors are predominantly found in regions of rapidly changing magnetic fields - around the calcification and close to tissue/air/bone interfaces, and surrounding vessels.

**Figure 6.**
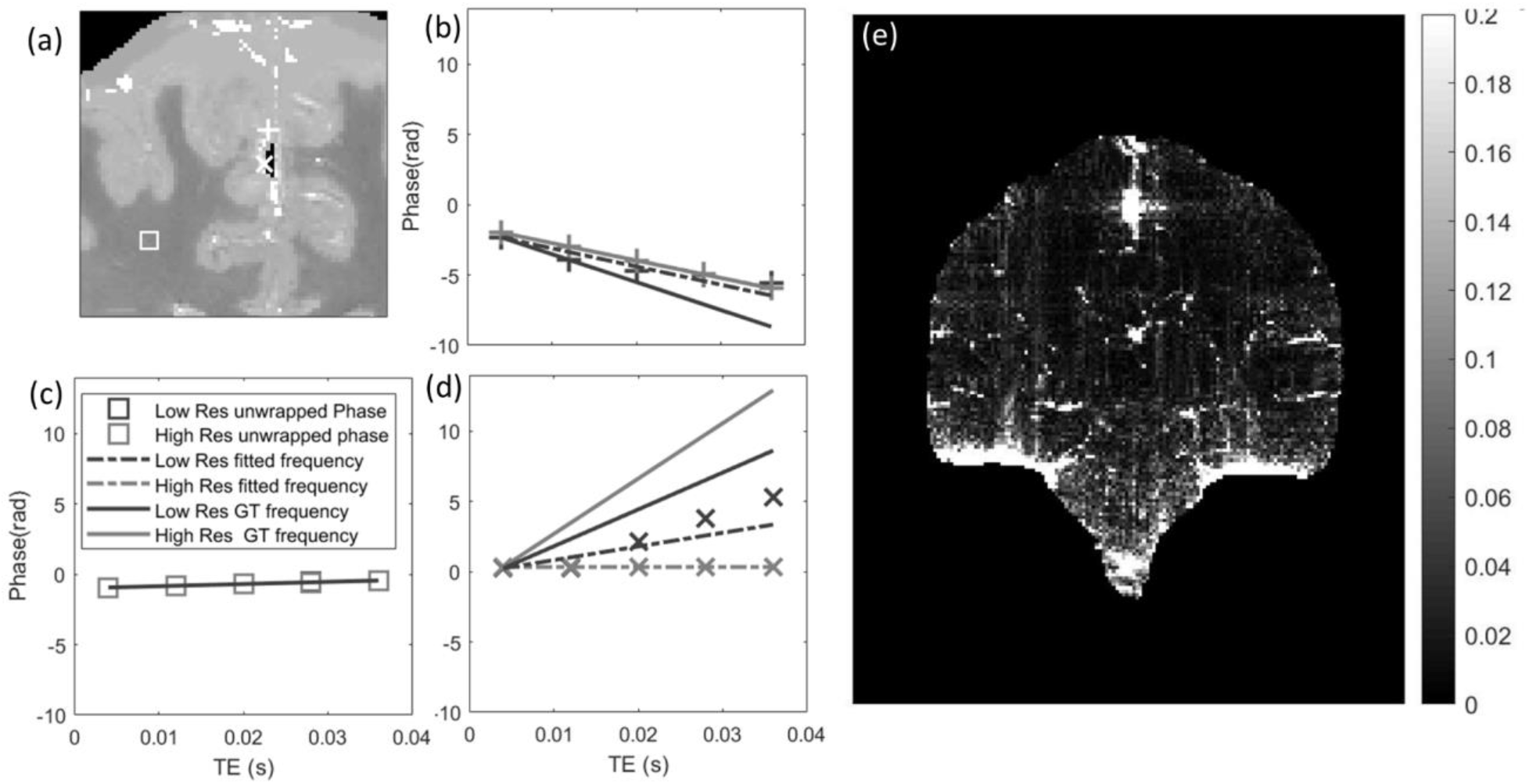
(a) Coronal view of the susceptibility phantom with three locations highlighted (square: region in the middle of white matter; cross and plus: two regions close to the calcification). (b,c,d) Plots of the unwrapped phase at the three locations (b – cross, c –square and d - plus) as a function of echo time. Unwrapped phase on the high-resolution data (light grey) and low-resolution data (dark grey) are shown using the respective markers, while dashed lines corresponds to the fitted frequency for each point at each resolution and the continuous line shows the ground frequency evolutions. (e) Mean squared difference map across echo times between fitted phase (dashed line in panel b-d) and measured phase (after unwrapping) on the coronal slice, highlighting tissue bone interfaces as well as regions surrounding the calcification.

### 3.2 Evaluation of QSM reconstructions

Figure 7 shows the reconstructions with minimum RMSE for the 4 algorithms tested. It can be seen that the direct methods (TKD and closed-form L2) still have some broad striking artifacts in regions surrounding both the calcification and deep gray matter regions while in the iterative methods these were reduced. It is interesting to note that the total variation (TV) regularized nature of FANSI clearly contributed to a better reconstruction of the superior cerebellar vein when compared to iLSQR. Please refer to the accompanying manuscript reporting the QSM 2.0 challenge results to see how these remaining artifacts were addressed by newer and more thoroughly optimized algorithms (23).

**Figure 7.**
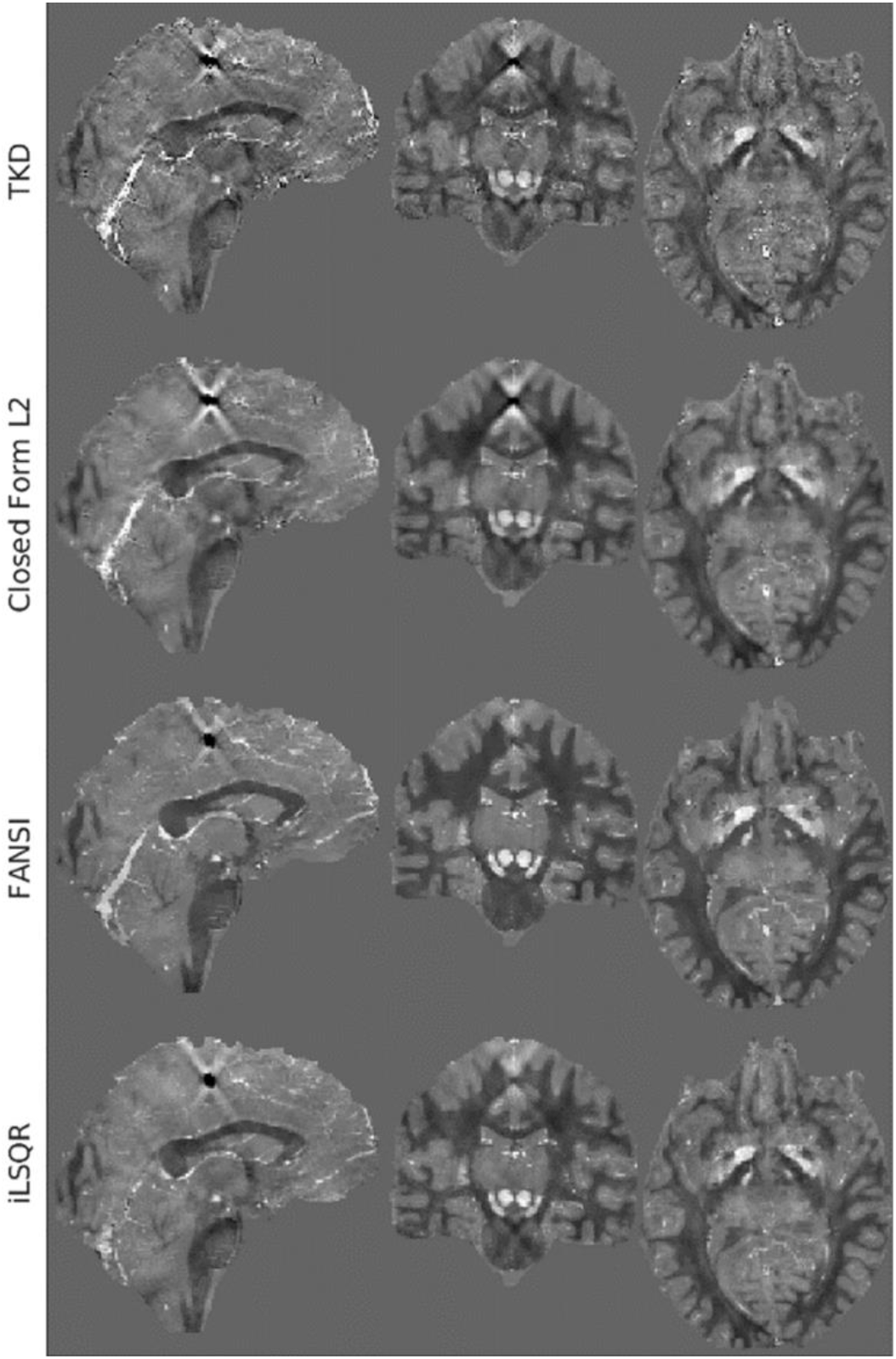
Examples of the optimum nRMSE reconstructions in three orthogonal planes obtained from the 1mm dataset based on SIM2 (see figure 2) with peak SNR 100 used for the QSM 2.0 challenge. The sagittal and coronal were chosen to cross the calcification region to highlight remaining streaking artifacts. The 4 different rows correspond to the 4 different reconstruction pipelines tested.

### 3.3 Comparison of simulations including microstructure to real data

Figure 8 shows an example of the originally acquired and the simulated magnitude and phase data, respectively. The magnitude data (first column) shows similar high resolution features at the matched echo time despite the acquisition protocols not being identical (the simulations only support multi-echo gradient echo acquisitions rather than MP2RAGEME as used for data acquisition). It can be visually observed that the simulated data suffers from reduced bias field inhomogeneity, this is a result of a bias field correction applied to the computed M_0_ map obtained from the MP2RAGEME. This choice was justified by two factors: (a) the magnitude bias field observed after SENSE (73) reconstruction does not reflect the local SNR but a mix of the volume and surface coil sensitivities as well as the transmit coil inhomogeneities; (b) we wanted to separate the physical simulation from the interaction with the hardware. In the non-background field phase data, the background field associated with nasal sinus and ear canals is weaker on the simulated data than on the measured data (as can be appreciate by the larger number of phase wraps on the latter). This can be attributed to two features, the imperfect bone air segmentation or an underestimation of the susceptibility differences between tissue and bone or air. The latter is also supported by observing on the third column (after background field removal) that the field surrounding the calcification is smaller in the simulations than that what is observed in the experimental data.

**Figure 8.**
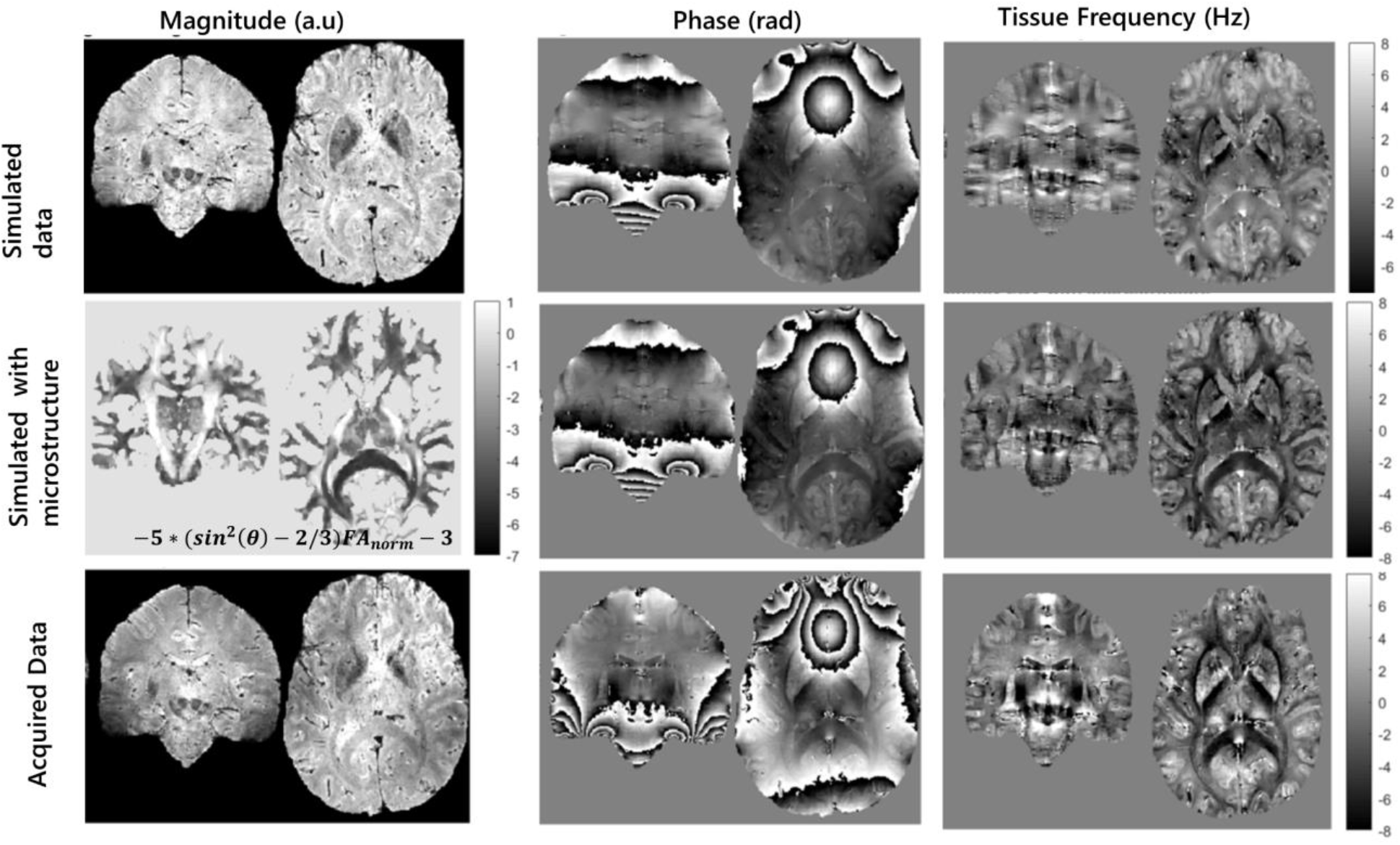
Shows the impact of adding a microstructure correction term to the simulated field map. First and second column show the simulated magnitude and phase data using whole brain susceptibility model at TE=20ms, while the third column shows the computed tissue frequency map (after brain masking and background field removal). First row shows the simulated data using Protocol 1 without the addition of the microstructure effects. The second row shows the impact of adding the microstructural effects as described in equation 7 (first column) both on the third echo time and the computed tissue frequency map. The third row shows, for visual comparison purposes, the experimental data acquired at 7T.

When comparing grey-white matter contrast on both the phase data (second column) and tissue frequency (third column), it can be see that the simulation better visually approximate the acquired data when the microstructural correction term (middle row) is added to the simulated data as expressed in Equation 7. This data can now be used to test how different reconstruction pipelines are biased due to microstructural effects. Note that, because our susceptibility phantom is based on reported values of susceptibility rather than this particular subject susceptibility values, no quantitative evaluation of the similarity can be performed.

## 4 Discussion

In this paper and the accompanying shared dataset and code (described in greater detail in the Supporting Information), we have presented and disseminated a realistic human brain phantom that can be used by both the QSM and the MRI communities to simulate multi-echo GRE data and evaluate QSM pipelines in a controlled manner.

The data and code provided allow users to:

- create new susceptibility phantoms, with different levels of spatial modulation of each compartment – this can be simply performed by changing the values presented on Table 1 which control both the mean value and spatial modulation present within each tissue;
- create realistic gradient echo multi-echo data at 7T and, to some extent, also at other fields (it should be noted unlike susceptibility, relaxation times are field dependent and their field dependence is tissue dependent).
- assess the impact of protocol changes (as well as nuisance factors such as RF phase, B_0_ shimming and noise) on the quality of the obtained QSM maps,
- assess the impact of changing some of the background field removal and QSM algorithm specific options while having a ground-truth to test it against.

This digital phantom will be important for QSM users and researchers deciding on acquisition protocols. Protocol considerations such as impact of the number of echo times and its range, as well as the degree of T_1_ weighting and resolution on the ability to accurately measure QSM in a given brain region can be quickly tested in this framework. This would allow for example to extend the analysis done by Karsa et al (74) on the effects of field of view and anisotropic voxels sizes, or the analysis by Biondetti et al on the effects of Laplacian based single echo vs multiple echoes techniques (75).

While the simulation framework is useful for optimizing protocols, the simulation in its current form is a static one. Consequently flow artifacts (which build up when large number of echoes are used), respiration related B_0_ fluctuations (76) and spatial distortions associated with the readout bandwidth are not considered. The latter two would be relatively straightforward to implement. Spatial distortions associated with different readouts can be obtained in computationally efficient manner by using our provided phantom data in, for example, freely available software packages such as JEMRIS, http://www.jemris.org/ (77). Wave Caipi (78) and 3D EPI (79,80) have been used for QSM, but not much research has been done to quantify the impact of their bluring or distortions on the performance of neither the background field removal nor the susceptibility maps. Respiration artifacts would simply require a library of respiration fields over time as acquired with field cameras (81). Flow artifacts would be more complex to simulate because the current vessel segmentation does not distinguish arteries and veins and it would be difficult to have a local flow velocity and pulsatility estimation.

The dipole model used in QSM assumes a sphere of Lorentz approximation (61), which does not hold true, particularly in white matter. The phantom released for the RC2 purposes, explicitly circumvented this limitation by ensuring perfect consistency with the QSM model in Eq. 4, i.e. no microstructural effects were present. With the released phantom dataset, we include diffusion data (both raw data on the 1.5mm space and its derivatives co-registered to the phantom space after distortion correction). These data can be used to either compute the frequency perturbation as shown in Figure 8, or the Hollow Cylinder Model can be used to explicitly introduce the echo-time varying perturbation similarly to what has been recently done in the context of myelin water imaging (22,82–85) and study the bias introduced by this effects on the reconstructed QSM maps. A critical challenge for more advanced modelling is the resolution of the diffusion acquisition, despite using state-of-the-art hardware and MR sequences. Here we have simply interpolated our 1.5mm DWI to the anatomical space and expect this to be sufficient to develop and validate QSM methods that account for microstructural effects in white matter (86).

QSM is gaining interest in the context of neurological disorders such as multiple sclerosis, PD and AD and other clinical applications such as hemorrhages or tumor imaging with iron oxide nanoparticles (87). For the latter applications, relaxation and susceptibility values in the form of, e.g., lesions in strategic locations can be easily added to the current phantom. Simulated data might be also relevant in the case of group studies of diseases, where differences in the deep grey matter nuclei were found (88,89) and the optimum QSM reconstruction parameters might improve the limits to detect those changes. Such question can be readily addressed with the digital phantom by changing the parameters of Eq. 1 for a given set of structures and test the QSM pipeline that can better quantify those changes.

## 5 Conclusion

The presented realistic and modular phantom aims to enable researchers to optimize reconstruction as well as acquisition parameters. As such, the phantom served as a ground-truth for the QSM RC2. Its modular design allows adding microstructure effects *a posteriori* (22) as well as the inclusion of new nuisances such as hemorrhages or fine vessels with realistic relaxation and susceptibility properties. We foresee that this brain model will be an important tool for the evaluation of various processes associated with QSM processing and interpretation.

## Supporting information

Support Information

## Data Availability Statement

The code used to create the phantom, as well as to generate the various simulations and figures described in this paper can be found on the data sharing collection: https://doi.org/10.34973/m20r-jt17

**Supporting Information Table S1:** Supplementary Information Table S1. Ad-hoc segmentation correction based on relaxometry values. Note that this corrections were only applied in regions where the magnetic field gradient, as computed from the multiecho data, was expected not to corrupt the R2* values;

## Supporting Information

The document has two sections. The first section outlines the main functionalities provided in the data sharing collection:

- creation of a realistic magnetic susceptibility phantom;
- simulation of a GRE data with a specific protocol;
- evaluation of a QSM reconstruction using provided phantom;
- adding microstructural information;

Finally it presents a detailed description of the file structure of the Data Sharing Collection associated with this publication.

The second section describes the methods and results of a preliminary evaluation of the QSM challenge dataset and the derived metrics as a mean to optimise 4 different reconstruction pipelines.

## Notes

https://doi.org/10.34973/m20r-jt17

### Competing Interest Statement

The authors have declared no competing interest.

### Summary of Updates

Changes were made in response to request from the MRM anonymous reviewers.

