## Supplementary material for "QSM Reconstruction Challenge 2.0: a realistic in silico head phantom for MRI data simulation and evaluation of susceptibility mapping procedures": Support Information

### 1 Data Sharing Collection

The code used to create the phantom described in this paper, the various simulations and figures can be found on the data sharing collection: https://doi.org/10.34973/m20r-jt17

In the following subsections some of the main functionalities provided are outlines:

- creation of a realistic magnetic susceptibility phantom;
- simulation of a GRE data with a specific protocol;
- evaluation of a QSM reconstruction using provided phantom;
- adding microstructural information;
- structure of Data Sharing Collection

#### 1.1 Creation of a realistic magnetic susceptibility phantom

Example code can be found on `MacroCreateSusceptibilityPhantom.m`.
The main function responsible by the creation of a new susceptibility phantom is `CreateOwnRealisticPhantom`, which takes as an input a matlab structure with the following fields:

- R1map_file – full path of R1 map in $s^{-1}$ in NIfTI format;
- R2starmap_file – full path of R2* map in $s^{-1}$ in NIfTI format;
- M0map_file – full path of M0 map in arbitrary units and in NIfTI format;
- Segmentation_file – full path of segmentation file in NIfTI format (each segmented tissue is associated with an integer);
- ChiModulation_file - matlab file containing the various parameters relating to Equation 1 and shown on Table 1 to create a susceptibility map – indices correspond to those used in the Segmentation nifty;
- rawField_file – full path of precomputed field map in NIfTI format;
- highGradMask_file – full path of binary mask in NIfTI format, defining regions where R2* values can and cannot be trusted, respectively;
- OutputChiModel_file – full path of address where to store the output susceptibility model in ppm units;

Furthermore, code in `MacroCreateSusceptibilityPhantom.m.` , creates Figures 2 and 3 of the current manuscript.

#### 1.2 Creation of simulated GRE data

Example code can be found in `MacroCreateSimulationdata.m`, with the main function being `CreateSimulationData` that takes as an input three Matlab structures. These Structures contain the information regarding the model parameters, the sequence parameters of the gradient echo sequence and the simulation parameters respectively.
The model parameters fields are:

- Chimap_file – location of magnetic susceptibility map in ppm units and NIfTI format;;
- R1map_file – location of R1 map in s-1 in NIfTI format;

- R2starmap_file – location of R2* map in s-1 in NIfTI format;;
- M0map_file – location of M0 map in arbitrary units in NIfTI format;
- Segmentation_file – location of segmentation file in NIfTI format;
- BrainMask_file = location of brain mask file in NIFTI format;

The sequence parameters fields define the GRE contrast and are : TR (repetition time, in secs), TE (echo time, in secs), FlipAngle (flip angle, degrees). Finally, the simulation parameters are:

- Res – spatial resolution of output in mm ;
- B0 and B0_dir – the main static magnetic field strength (in Tesla) and its orientation with respect to the head model (if along $B_0$ is along z, B0_dir=[0 0 1], if if along $B_0$ is along x, B0_dir=[1 0 0]);
- PhaseOffset - multiplier term of a quadratic phase over the brain aimed at simulating the transceive phase (0 : no phase offset, 1 : a π phase difference inside the brain mask ;
- Shimm - boolean variable to activate B0-shimming with second-order spherical harmonics (0 : no B0 shimms is applied, 1 : 2nd order shimms are computed inside the brain mask ; works only when doing the simulations with the whole head model)
- Output_dir - Directory where the simulation results will be saved

Running the function `CreateSimulationData` creates two directories in Output_dir, "SimulatedHR" (where the native resolution simulation data is stored) and "Simulated_1p0mm" (where the downsampled data is stored, in the example above 1.0mm isotropic). For each sequence protocol two simulations are performed: one with the full head susceptibility model, and for the brain masked susceptibility model [Eq. 5].

Downsampled simulation results are stored as 3D or 4D (depending on the number of echo times) using the following file naming convention :
Data[Extension]_magn.nii.gz and
Data[Extension]_phase.nii.gz,
where "Extension" is either empty or ‘BrainExtracted’.
Additionally, the downsampled directory contains the new brain mask (Brain.nii.gz), the downsampled Segmented NIfTI (FinalSegment.nii.gz) and the downsampled susceptibility maps (Chi_crop.nii.gz).

Finally, the script `MacroCreateSimulationdata.m` reproduces the images in Figure 4.

#### 1.3 QSM pipeline evaluation

The script `MacroCreateQSMpipelineAndCompareHighResLowRes.m` creates the images in figures 5 and 6 of the current manuscript. Furthermore it illustrates how to create a data structure for perform a QSM reconstruction using SEPIA (sepia-documentation.readthedocs.io/) and evaluate the output using the various metrics used in RC2 and introduced in the Methods section of this manuscript.

The function `EvaluateRecon_ChallengeFinalMetricsRelease` takes as input the reconstruction file name and a structure with four fields:

- Segment - Segmentation file at a matched resolution (output of `CreateSimulationData`);
- chi_crop - Ground truth susceptibility map file obtained by k-space cropping (output of `CreateSimulationData`);
- maskEroded - brain mask where evaluation is to be performed;
- CalcMoment – calculated moment of the calcification performed on the high resolution susceptibility map;

The output of the function contains the metrics described in methods section.

#### 1.4 Comparison of simulation data to real data

The script `MacroAddingMicrostructure` contains the implementation of the code needed to generate Figure 8 using the already pre-processed DWI. The addition of the microstructure data is as described in Eq. 7.

#### 1.5 Structure of the Data Sharing Collection

Description of directory structure and files provided in the data sharing collection:

```
│   MacroAddingMicrostructure.m
```

*% example of how to add microstructure effects on the computed frequency shit maps – Figure 9*

```
│   MacrocreateQSMpipelineAndCompareHighResLowRes.m
```

*% example of how to use the simulated data to create a QSM pipeline in the high or low resolution data*

```
│   MacroCreateSimulationData.m
```

*% example of how to use the provided code (along with relaxation and susceptibility maps) to generate different complex datasets – with and without effects from background (air/bone/muscle/shims) fields and residual fields (RF)*

```
│   MacroCreateSusceptibiltyPhantom.m
```

*% example code on how to generate a susceptibility map using the relaxation rate maps and a table describing the mean values and modulation values within each segmented region*

```
│   MacroLoopReconstructionsOfPhantoms.m
```

*% evaluation of reconstructions using different supported methods - Figures 7 & 8*

```
│   MacroProcessInvivoData.m
```

*% example of how to process the invivo data on the data/raw directory into a QSM map*

```
│
├───data
```

*Directory with all input data in the native resolution*

```
│   ├───chimodel
```

*Generated Chi models and the paramters used to derive them*

```
│   │       Chi.nii.gz
│   │       ChiModelMIX.nii             -QSM Challenge Sim2
│   │       ChiModelMIX_noCalc.nii      -QSM Challenge Sim1
│   │       FinalLabelSuscep2.mat
```

```
│       │      FinalLabelSuscepCalcificationFree.mat
│       │      paramaters.mat
│       │      paramatersCalcificationFree.mat
│       │      ProportinalityTermsForSusceptibility.mat
│       │
│       ├───maps
```

*Directory with relevant relaxation rate maps*

```
│       │      M0.nii.gz
│       │      R1.nii.gz
│       │      R2star.nii.gz
│       │
│       ├───masks
```

*Masks associated with the digital phantom*

```
│       │      BrainMask.nii.gz
│       │      BrainMaskBrainExtracted.nii.gz
│       │      highgrad.nii.gz
│       │      SegmentedModel.nii.gz
│       │
│       └───raw
```

*Directory containing the raw data used to compute the relaxation maps*

*Derived data from the 3T data DWI (fractional anisotropy, FA, and main tensor Orientation, V1) are given already on the high resolution imaging space*

```
│           │  MP2RAGEME_brain.nii.gz
│           │  MP2RAGEME_INV1.nii
│           │  MP2RAGEME_INV1ph.nii
│           │  MP2RAGEME_INV2.nii
│           │  MP2RAGEME_INV2ph.nii
│           │  MP2RAGEME_INV2_e2.nii
│           │  MP2RAGEME_INV2_e2ph.nii
│           │  MP2RAGEME_INV2_e3.nii
│           │  MP2RAGEME_INV2_e3ph.nii
│           │  MP2RAGEME_INV2_e4.nii
│           │  MP2RAGEME_INV2_e4ph.nii
│           │  MP2RAGEME_total-field_ppm_head_r2s-header.nii.gz
│           │
│           └───sepia
```

*Processed MP2RAGEME data for the propose of QSM computation and comparison with simulated data*

```
│           └───3T
│               └───Derived_HighResSpace
│                       V1.nii.gz
│                       FA.nii.gz
```

*Processed diffusion data corregistered to the High res 7T data*

```
│
```

```
├───func
```

*Folder with relevant functions used in the production of phantoms and simulation of data*

```
│    │   CreateOwnRealisticPhantom.m
│    │   CreateSimulatedData.m
│    │
│    │
│    ├───evaluation
```

*Evaluation scripts*

```
│    └───utils
│         ├───nii
│         ├───reisert-unring-ddd43da65219
Unringing
│         └───WaveletDenoising
├───ManuscriptFigures
│    ├───BackgroundFieldRemoval
│    │    └───unwrapping
│    ├───BackgroundFieldRemovalBrainExtracted
│    │    └───unwrapping
│    ├───BackgroundFieldRemovalHR
│    │    └───unwrapping
│    └───BackgroundFieldRemovalHRBrainExtracted
│         └───unwrapping
└───Simdata
```

*Folder containing Simulated data for various MR protocols*

```
     │   SimSeqParams.mat
     ├───DGMprotocol
```

*Folder with simulated data with protocol aimed at measuring Deep gray matter susceptibility*

```
     │    │   SimulationParameters.mat
     │    ├───SimulatedHR
```

*High Resolution simulation using the parameters described in SimulationParameters.mat*

*It contains the 4D magnitude and phase data at the different echo times with the simulations performed using whole head or brain tissue only sources of susceptibility*

```
     │    │        Data_magn.nii.gz
     │    │        Data_phase.nii.gz
     │    │        Data BrainExtracted_magn.nii.gz
     │    │        Data BrainExtracted_phase.nii.gz
     │    └───Simulated_1p0mm
     │
```

*Folder containing the down-sampled simulated data of the above mentioned protocol. Additionally the brain Segmentation model, brain masks and susceptibility were equally down-sampled to the desired resolution so that a ground truth at the target resolution exists. You can find datasets with and without the extension 'BrainExtract', it refers to the cases where brain tissues susceptibility only or whole head susceptibilities were used for the simulation*

```
│   │       Brain.nii.gz
│   │       BrainBrainExtracted.nii.gz
│   │       Chi_crop.nii.gz
│   │       Chi_cropBrainExtracted.nii.gz
│   │       Chi_interp.nii.gz
│   │       Chi_interpBrainExtracted.nii.gz
│   │       FinalSegment.nii.gz
│   │       FinalSegmentBrainExtracted.nii.gz
│   │       Data_magn.nii.gz
│   │       Data_phase.nii.gz
│   │       Data BrainExtracted_magn.nii.gz
│   │       Data BrainExtracted_phase.nii.gz
│
└───GMWMprotocol
```

*Same substructure as above but for a protocol aimed at differentiating grey and white matter*

```
│
└───ChallengeProtocol
```

*Same substructure as above but now using the protocol used in Challenge2.0 -*

### 2 QSM pipeline optimization for QSM challenge setting

#### 2.1 QSM reconstruction optimization

To demonstrate the applicability of the current framework for the QSM challenge or for QSM reconstruction optimization purposes, we performed a simulation with only local fields (see Eq. 5). The protocol used was that of the QSM challenge (TR = 50ms; $TE_{1/2/3/4}$= 4/12/20/28ms; α=15). Pre-processing was the same as mentioned on the previous subsection, with the exception of LBV, which was not required. QSM reconstruction using TKD (63), closed form L2 (64), FANSI (65) and iLSQR (66), respectively, with each of the QSM reconstructions being run 20 times with varying regularization parameters:

- TKD: the k-space threshold was varied from 0.02 to 0.4 in linear steps;

- Closed form L2: the regularization parameter was varied in 20 logarithmic steps 0.01 to 100;

- FANSI TV: the regularization parameter as varied from $10^{-6}$ to 0.1 in logarithmic steps, remaining parameters were $\mu_1$=0.005, $\mu_2$=1 and 50 iterations; note that the cost function used in FANSI is the same as that first introduced in MEDI (67) but with a faster solver;

- iLSQR: the regularization term was varied from $10^{-3}$ to 1 in logarithmic steps, had a tolerance of 0.01 and was allowed 50 iterations.

The reconstructions were evaluated in respect to the ground-truth using various metrics tested for the purpose of the QSM challenge and described in Table 1 of the main manuscript.

The process was repeated with different simulated datasets:

- 1mm and 0.64mm isotropic resolution
- With infinite SNR (no noise added) or peak SNR of 100

#### 2.2 Evaluation of QSM reconstructions

Figure S1a shows the nRMSE values associated with various reconstruction methodologies as a function of the regularization value for the high-resolution phantom in the absence of noise. Figure S1b shows the more realistic case where the images have been down-sampled to 1mm isotropic and noise was added to the data with a peak SNR of 100. It can be seen from a shift of the individual curves to the right that the more realistic phantom required stronger regularization than the perfect high-resolution phantom, which can be attributed to the phase noise and inconsistencies associated with the down-sampling process. The FANSI algorithm achieved a lower nRMSE in respect to the down-sampled phantom than on respect to the original phantom (36 vs 43), the same being true, to a smaller extent, for the closed form L2 methods. Although one would expect the error to always increase, as observed in Fig. 5. Care should be taken when comparing the nRMSE across different resolutions as the normalization factor changes, another potential source of confounds is the down sampling and unringing strategy used for the susceptibility map that could introduce some spatial smoothness similar to that enforced both by FANSI and closed form L2. Figure 7c shows the reconstructions with minimum RMSE of the plots in Fig. 7b for the different methods tested. It can be seen that the direct methods (TKD and closed-form L2) still have some broad striking artifacts in regions surrounding both the calcification and deep gray matter regions while in the iterative methods these were reduced. It is interesting to note that the total variation (TV) regularized nature of FANSI clearly contributed to a better reconstruction of the superior cerebellar vein when compared to iLSQR. Figures S1d and S1e show the performance of iLSQR and FANSI respectively for the different resolutions and noise levels. It is interesting to observe that, for both methods, both the down sampling and the addition of noise result in an increase of the optimum regularization value associated with the additional phase inconsistencies.

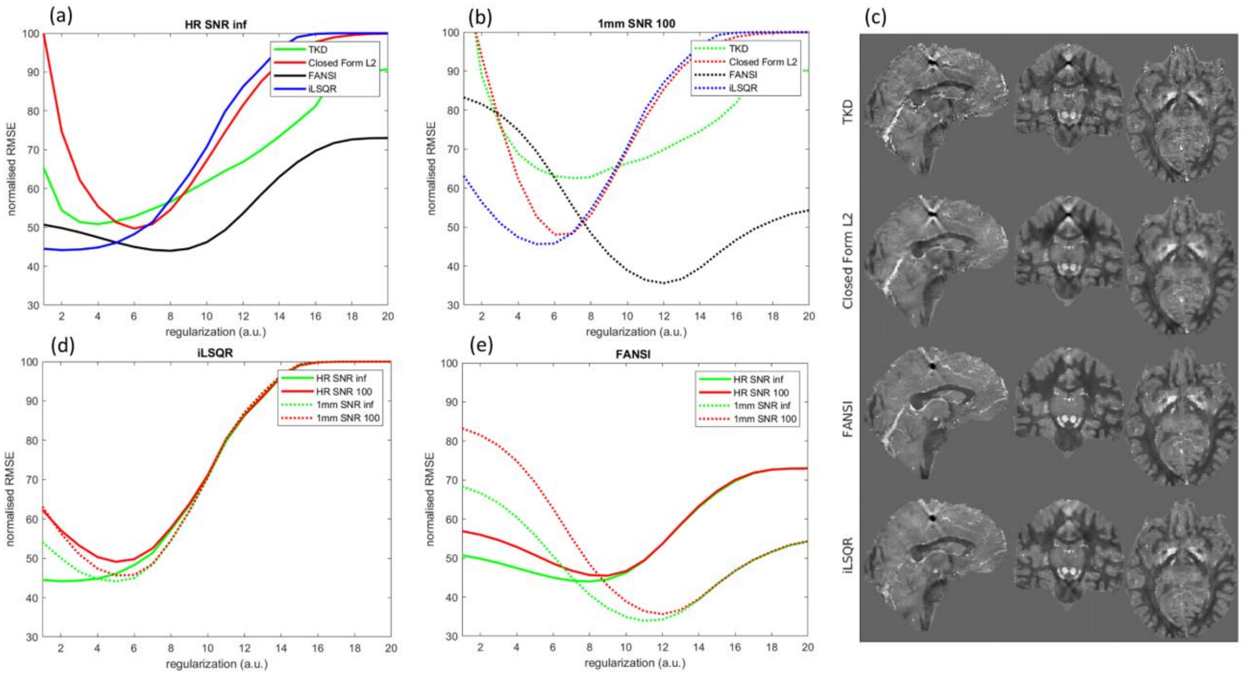

**Figure S1** Shows various plots of the normalized RMSE as a functions of the regularization value (a,b,d,e). Plots (a) and (b) show the numerical performance of 4 different QSM algorithms when the input data was the high resolution simulated data with no additional noise or the down-sampled 1mm data with added noise resulting in an image peak SNR of 100. Plots (d) and (e) show the performance of iLSQR and FANSI respectively for different types of input data: high resolution and 1mm data shown in solid and dashed lines respectively; without and with added noise shown in green and red. Panel (c) shows examples of the optimum nRMSE reconstructions of the 1mm dataset with peak SNR 100 using the 4 reconstruction algorithms tested in the manuscript.

Finally, more specific metrics were evaluated. These included RMSE in different tissues (Fig. S2a), calcification moment calculation and linearity susceptibility estimations in white matter. It is interesting to observe that, for each method, the optimum regularization values for the different regions remains mainly unchanged, with the minimum of each curve corresponding to either the same regularization factor or minimal deviations when compared to the broadness of the curves. Interestingly, when comparing the dependence of the calculation of the calcification moment (measured as -49.8 ppm on the high resolution phantom) as a function of the regularization parameter, it can be observed that the two evaluated methods have different optimum regularizations when compared to their optimum RMSE settings (iLSQR and FANSI needed a lower and a higher regularization factors, respectively). Figure S2c, shows the expected behavior where the greatest precision of the accuracy of deep gray matter susceptibility estimation (as measured by the slope between reconstructed average QSM value per deep gray matter structure and ground-truth) was obtained using the smallest amount of regularization (29,74).

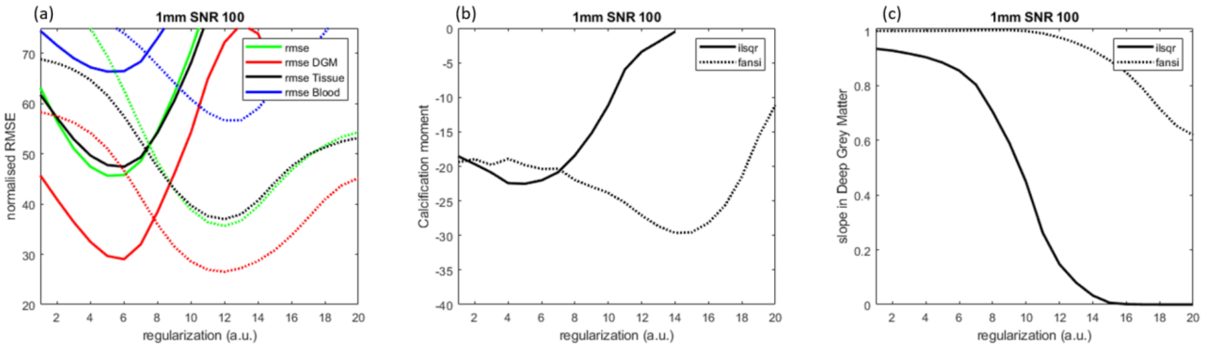

**Figure S2** (a-c) Plots of the various metrics as a function of the used regularization value for the iLSQR (continuous line) and FANSI (dashed line) algorithms. The different plots show: (a) normalized RMSE reconstructions metrics, (b) calcification metric and (c) linearity of the susceptibility values measured in deep grey matter in respect to ground truth.

### 3 Suplemental Information Table S1

| Initial tissue segmentation | R1(mHz) criteria | R2*(mHz) criteria | Final tissue segmentation |
|---|---|---|---|
| CSF | -<br>R1>=250 | -<br>R2 > 70 | CSF<br>Blood |
| Grey matter | -<br>R1<330<br>R1>=330<br>R1 > 750 | -<br>-<br>R2 >= 100<br>R2 < 100 | Grey matter<br>CSF<br>Blood<br>WM |
| Caudate | -<br>R1<330<br>R1>=330<br>R1 > 750 | -<br>-<br>R2 >= 100<br>R2 < 100 | Caudate<br>CSF<br>Blood<br>WM |
| Putamen | -<br>R1<330<br>R1>=330<br>R1 > 750 | -<br>-<br>R2 >= 100<br>R2 < 100 | Putamen<br>CSF<br>Blood<br>WM |
| Thalamus | -<br>R1<330 | -<br>- | Thalamus<br>CSF |

| | | | |
|---|---|---|---|
| | R1>=330<br>R1 > 750 | R2 >= 100<br>R2 < 100 | Blood<br>WM |
| White matter | -<br>R1<330<br>R1>=330 | -<br>-<br>R2>100 | White matter<br>CSF<br>Blood |
| Blood (derived from Frangi filter maximum Diameter 6 voxels) | | R2>75 | Blood |

**Supplementary Information Table S1.** Ad-hoc segmentation correction based on relaxometry values. Note that this corrections were only applied in regions where the magnetic field gradient, as computed from the multiecho data, was expected not to corrupt the R2* values
